# Gene Sets Analysis using Network Patterns

**DOI:** 10.1101/629816

**Authors:** Gregory Linkowski, Charles Blatti, Krishna Kalari, Saurabh Sinha, Shobha Vasudevan

**Author notes:** To whom correspondence should be addressed. Email: Shobha Vasudevan,. Correspondence may also be addressed to Saurabh Sinha,. These authors contributed equally to this work (Gregory Linkowski, Charles Blatti).

## Abstract

High throughput assays allow researchers to identify sets of genes related to experimental conditions or phenotypes of interest. These gene sets are frequently subjected to functional interpretation using databases of gene annotations. Recent approaches have extended this approach to also consider networks of gene-gene relationships and interactions when attempting to characterize properties of a gene set. We present here a supervised learning algorithm for gene set analysis, called ‘GeneSet MAPR’, that for the first time explicitly considers the patterns of direct as well as indirect relationships present in the network to quantify gene-gene similarities and then report shared properties of the gene set. Our extensive evaluations show that GeneSet MAPR performs better than other network-based methods for the task of identifying genes related to a given gene set, enabling more reliable functional characterizations of the gene set. When applied to the set of response-associated genes from a triple negative breast cancer study, GeneSet MAPR uncovers gene families such as claudins, kallikreins, and collagen type alpha chains related to patient’s response to treatment, and which are not uncovered with traditional analysis.

## INTRODUCTION

Gene set analysis is a mainstay of functional genomics studies today. High-throughput experiments allow researchers to identify a set of genes related by a common profile, such as differential expression between experimental conditions, altered epigenomic states in regulatory regions, or signatures of selection in their coding sequences. It is then common to seek other properties (e.g., Gene Ontology terms and pathways) of relevance to the gene set so as to make connections between those properties and the original defining property of the gene set. This task is usually approached by testing if the gene set is significantly enriched for genes annotated with a property, iterating over many, even thousands of properties; tools such as DAVID (1) and GSEA (2) offer such enrichment analysis through convenient interfaces, and have been widely adopted. We refer to this task as ‘gene set characterization’ (3). A second line of research into gene set analysis seeks to identify genes most related to a given gene set. For instance, numerous methods for predicting functions of poorly annotated genes begin with a set of genes in a Gene Ontology (GO) category and use pre-determined gene-gene networks to identify additional genes that might share that function (4–6). These methods utilize some version of the ‘guilt-by-association’ principle: if a gene is directly or indirectly linked via the network to a gene with a specific function then it might share the latter’s function. We refer to this second type of gene set analysis as ‘gene set membership prediction’ (GSMP), recognizing that the approach may be used to identify related genes of a gene set with any common property, not only GO annotations.

In this work, we draw inspiration from both lines of research outlined above to develop a new framework for gene set analysis. In particular, we develop a new network-based method for the GSMP task, demonstrate its advantages over state-of-the-art methods, and then use its predictions to perform the gene set characterization task. We believe that the two tasks are related and that an improvement in performing the GSMP task can lead to better, or at least novel, functional characterizations of a gene set. The rationale for this is the following: Gene set characterization as it is performed today amounts to identifying properties that distinguish genes in the given set from all other genes. However the full complement of genes may include genes that are closely related to the given set, and share the latter’s properties. If we could identify these closely related genes up front then we could seek properties that are common to the given gene set *and* these closely related genes, and not shared by the remaining, less related genes. That is, the contrast between the gene set and ‘all other genes’ can improve when we extend the given gene set to include its most closely related genes. Since it is difficult to systematically benchmark a gene set characterization method, we support the above premise by applying the new method to a clinically significant gene set and noting that it identifies biologically reasonable properties that would have been missed by traditional methods. On the other hand, the GSMP task itself is amenable to systematic benchmarking, since the evaluator may hide a part of a biologically coherent gene set and test if one GSMP algorithm can uncover those hidden genes better than other algorithms. We adopt this systematic approach to test GSMP performance of Geneset MAPR in comparison with other state-of-the-art algorithms on many gene sets and find clear and consistent improvements.

Methodologically, the distinguishing aspect of Geneset MAPR is its ability to exploit multiple gene networks simultaneously when characterizing gene-gene similarity and scoring any gene for its relationship to a given gene set. Handling of multiple gene networks together is an example of the ‘heterogeneous network mining’ problem, which presents daunting technical challenges in many domains of information science (7,8). (We will use the term ‘heterogeneous network’ here to refer to a network where multiple gene networks, each defined by a different type of relationship, are overlaid to create a single gene network with multiple edge types.) Heterogeneous network mining is germane in the genomics context because gene networks can be reconstructed from a variety of sources, e.g., protein-protein interaction data, homology information, genetic or regulatory interactions, etc. which carry complementary information about gene similarities that may bear upon a ‘guilt-by-association’ analysis. Despite this, most approaches to such an analysis make the simplifying assumption that the edges are of the same type (9–11), e.g., only protein-protein interactions, with a few notable exceptions. One approach is to combine different edge types in the heterogeneous gene network to form a homogeneous network prior to gene set analysis, as in the popular tool GeneMANIA (6). Another approach, exemplified by our previous algorithm called DRaWR (3), retains heterogeneous edges and performs a ‘random walks with restarts’ to quantify gene-gene connectivity. However, neither method treats different types of connectivity among genes as being of potentially different value to the GSMP task. (Here different ‘types of connectivity’ may refer to, for example, when two genes are connected because both exhibit protein-protein interaction with a third gene, or when one gene is homologous to and the other has a common GO annotation with a third gene). This is the key conceptual gap in current network-based gene set analysis methods that we address in the present work.

To address the above problem, Geneset MAPR (or ‘MAPR’, for brevity) quantifies network connectivity between genes while preserving information about *types* of connectivity. Central to the method is the ‘meta path’, a concept from graph mining that characterizes paths connecting a pair of nodes based on the sequence of edge types appearing in the path (12). MAPR uses meta paths to track different types of connectivity patterns in the heterogeneous gene network, and develop them to characterize the underlying relationships in a given gene set. Our choice of meta paths as a network analysis entity stems from the hypothesis that if different edge types in a heterogeneous network are treated as different, the analysis may provide a novel, valuable perspective on gene-gene relationships. A second major feature of MAPR is its use of supervised machine learning (‘classification’) for the GSMP task. We note that GSMP is at its heart a classification task – a set of genes belonging to a class (the gene set) is made available and the goal is to predict additional members of the class. MAPR embodies this perspective, casting the GSMP task as a classification problem, learning to predict membership in the gene set using a comprehensive description of each gene in terms of its meta path-based network connectivity patterns.

We show by cross-validation tests that Geneset MAPR achieves consistently better GSMP accuracy compared to GeneMANIA, DRaWR and other baselines, over nearly 600 different gene sets from 13 different compendia. The network edge types most useful for the task are automatically learned and vary from one compendium to another; gene-gene relationships based on text-mining proved to be much more useful for certain types of gene sets compared to others, while information from Gene Ontology annotations was consistently useful across most gene sets. We noted that gene sets comprising genes with better annotations in knowledge-bases or genes related to more widely researched topics such as cancer and drugs tend to be more predictable in the GSMP setting. Finally, we applied the new GSMP tool to a gene set associated with poor response to chemotherapy in breast cancer and identified three gene families – claudins, kallikreins, and collagens – that were not conspicuous in the original gene set but were highly ranked among the most closely related genes of the set. Prior work associates these gene families with poor prognosis, and thus lends credence to the new insights gleaned by our analysis about key genes in the target gene set.

## RESULTS

### Outline of Geneset MAPR, a new method for gene set membership prediction

The central task of interest to us is ‘gene set prediction’ (GSMP): given a gene set, find other genes ‘highly related’ to that gene set. To make the problem more objectively defined, we consider the following formulation of GSMP: *given a random subset of a gene set, predict the remaining genes of that set, from among all other genes*. To pose the GSMP task one must also specify the information that is available in performing the task. Here, we consider one or more compendia of prior knowledge about genes as the available information. Each compendium may include information about (i) genes annotated with specific properties, such as Gene Ontology biological processes, or (ii) pairs of genes related by a specific relationship, such as direct protein-protein interaction. We utilized 11 such compendia in the public domain (13–18), listed in Table 1. A GSMP method is required to make its predictions based on the knowledge stored in one or more of these compendia.

**Table 1.**
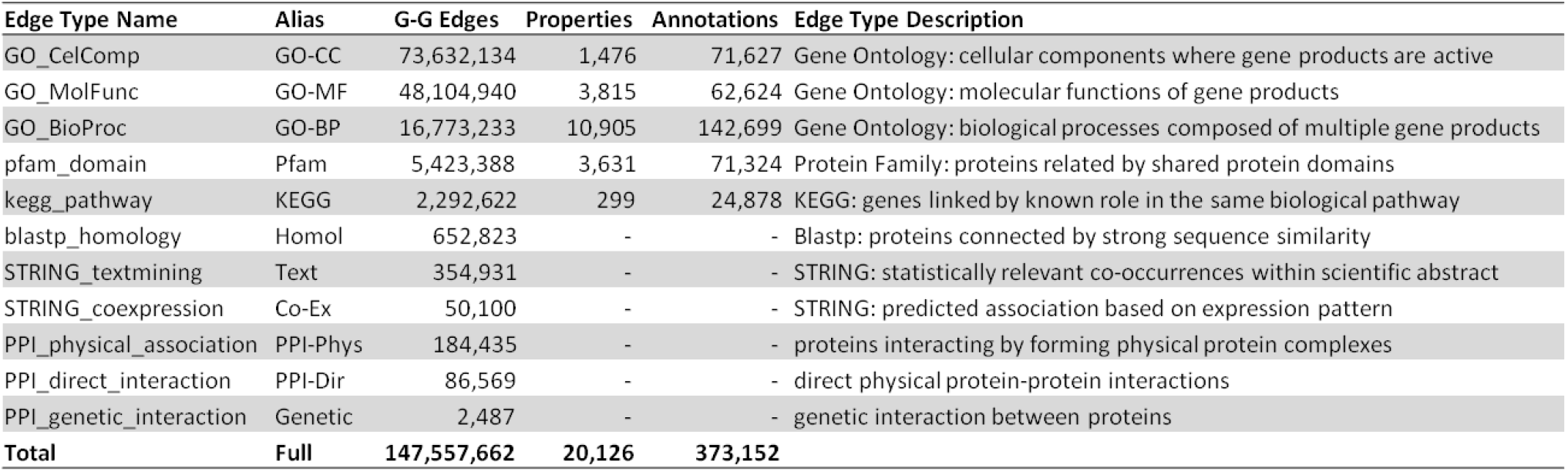
Heterogeneous Compendia of Public Knowledge. Each row shows the number of gene-gene edges of a given type derived from each different compendium of prior knowledge, also providing a type alias and description. For edge types that were derived from property-gene annotations, the number of distinct property nodes and total annotations are also included.

#### Compendia of prior knowledge as a heterogeneous network

We propose a novel method for GSMP, called ‘Gene Set Meta path Analysis for Pattern Regression’ or simply ‘GeneSet MAPR’ or ‘MAPR’ for short. It utilizes the available/provided compendia of prior knowledge by representing them as a heterogeneous network, i.e., a network where nodes represent genes and different types of edges represent prior knowledge from different compendia. It then solves the GSMP problem by defining the similarity of each gene to the given gene set through patterns of gene-gene connectivity in the heterogeneous network. We first note that each compendium may be represented as a homogeneous network. A compendium of gene-gene relationships (of a specific type such as protein-protein interaction) can be represented by an all-gene network, with edges recording those relationships. For a compendium of gene-property associations the information may be recorded as edges in a bipartite graph with gene and property nodes; MAPR transforms such a network into one that only has gene nodes, where a pair of nodes is connected by an edge if both genes are associated with the same property (Figure 1A,B). Thus, every compendium provided as prior knowledge is represented by a different homogeneous network involving genes, and a heterogeneous network is created by overlaying all of these networks (Figure 1C). The network edges may be weighted, if the compendia that they were derived from provide confidence values for relationships. All edges in the network are undirected, i.e., they represent symmetric relationships.

**Figure 1.**
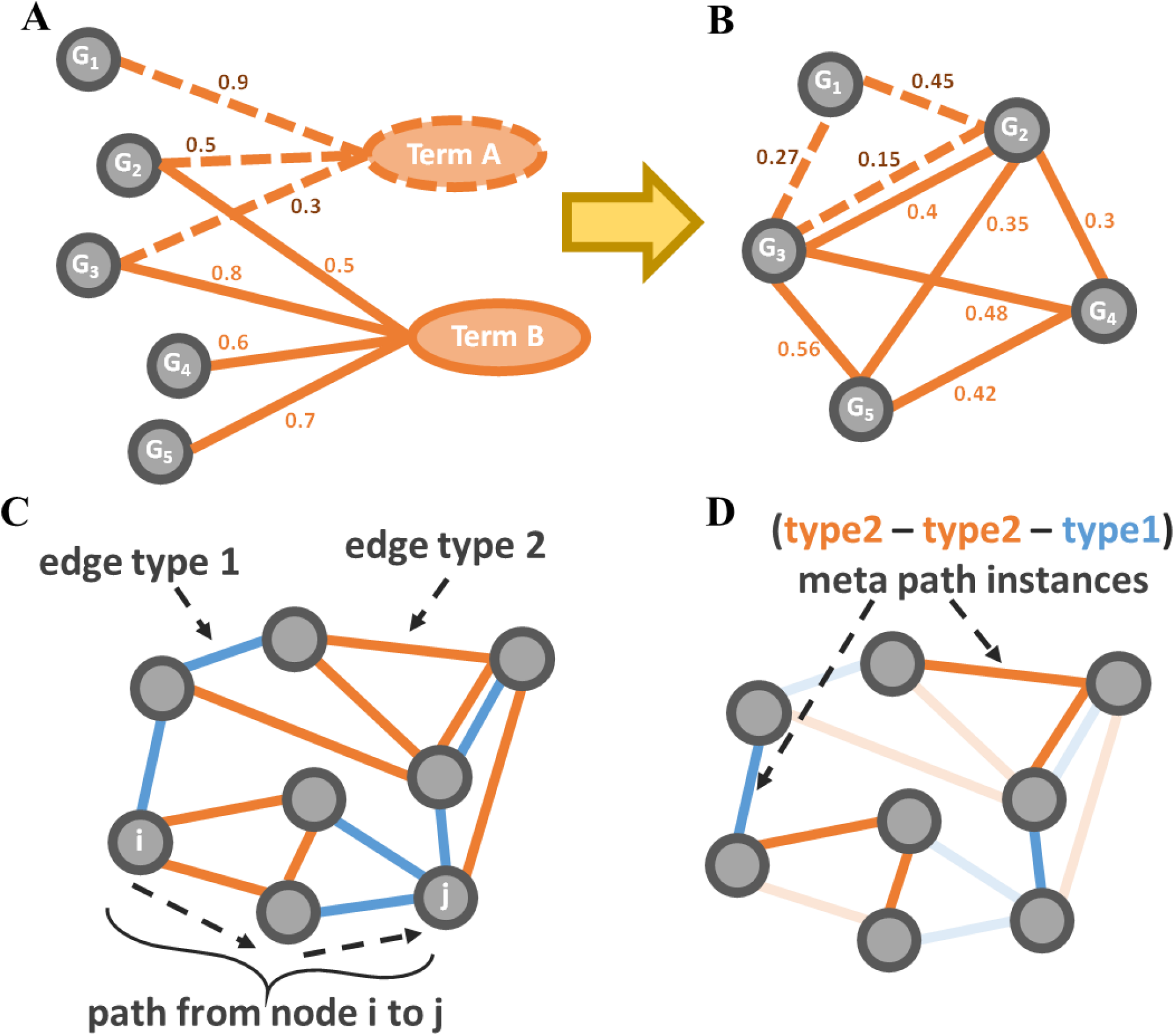
Understanding MAPR Meta Paths. A) A set of gene annotations can be represented as a bipartite graph, where genes (gray nodes) and annotation terms (colored nodes) can be connect with edges representing the annotation of the gene for the term and a weight indicating the confidence of that annotation. B) These bipartite graphs are converted to gene-gene networks in MAPR where the new gene-gene edges score the fact that connected genes share specific annotation relationship(s). C) Example heterogeneous network is shown with gene nodes (gray circles) and two different types of gene-gene edge relationships (orange and blue). A single path in this network is indicated with dotted arrows and a curly brace. D) In the same network, two instances of a specific meta path, *m*, are highlighted. The start and end nodes of the specified paths are said to be “connected along meta path *m*”.

#### Meta paths as connectivity patterns in heterogeneous network

MAPR examines the resulting network in terms of its connectivity patterns as defined by paths and ‘meta paths’. A path is a contiguous sequence of edges where each pair of successive edges is incident to a common node (Figure 1C), and is an intuitive way to describe the connectivity between two nodes. In a heterogeneous network, a meta path is an ordered sequence of edge types, such that any number of individual paths in the network may follow that sequence of edge types (12) (Figure 1D). Thus, meta paths are a means to categorize different paths in a heterogeneous network. They allow us to go beyond merely talking about strengths of connectivity between two nodes in the network (e.g., based on the counts of paths between them) to *types* of connectivity between them.

#### MAPR as a classifier using connectivity features

We briefly describe how MAPR, given a gene set *G*, ranks all genes (including genes in *G*) for similarity to *G*, thereby solving the GSMP problem. (See Figure 2 and **Methods** for details.) It first quantifies the *strength* of connectivity (henceforth, ‘linkage’) between each pair of genes in the network, separately for each *type* of connectivity (meta path, symbolized by ‘*m*’). The linkage (between two genes) takes into account the number of paths matching that meta path as well as the weights (if any) of the edges constituting those paths. A total linkage via meta path *m* is then computed between each gene and the query gene set *G*, by aggregating the gene’s linkage to each gene in *G*. Each gene is thus assigned a ‘feature vector’ where each dimension represents its total linkage to *G* via a different meta path. Genes in *G* are also assigned such feature vectors. Next, MAPR trains an ensemble of LASSO (19) regression models using these feature vectors, treating genes in *G* as the positive set and equally many randomly chosen genes (not in *G*) as the negative set. (Each random choice of negative set yields one model in the ensemble.) This ensemble of models is used to predict the gene’s membership in set *G*. Note that genes in the positive set *G* are likely to receive high scores, but are not guaranteed to be the highest scoring genes.

**Figure 2.**
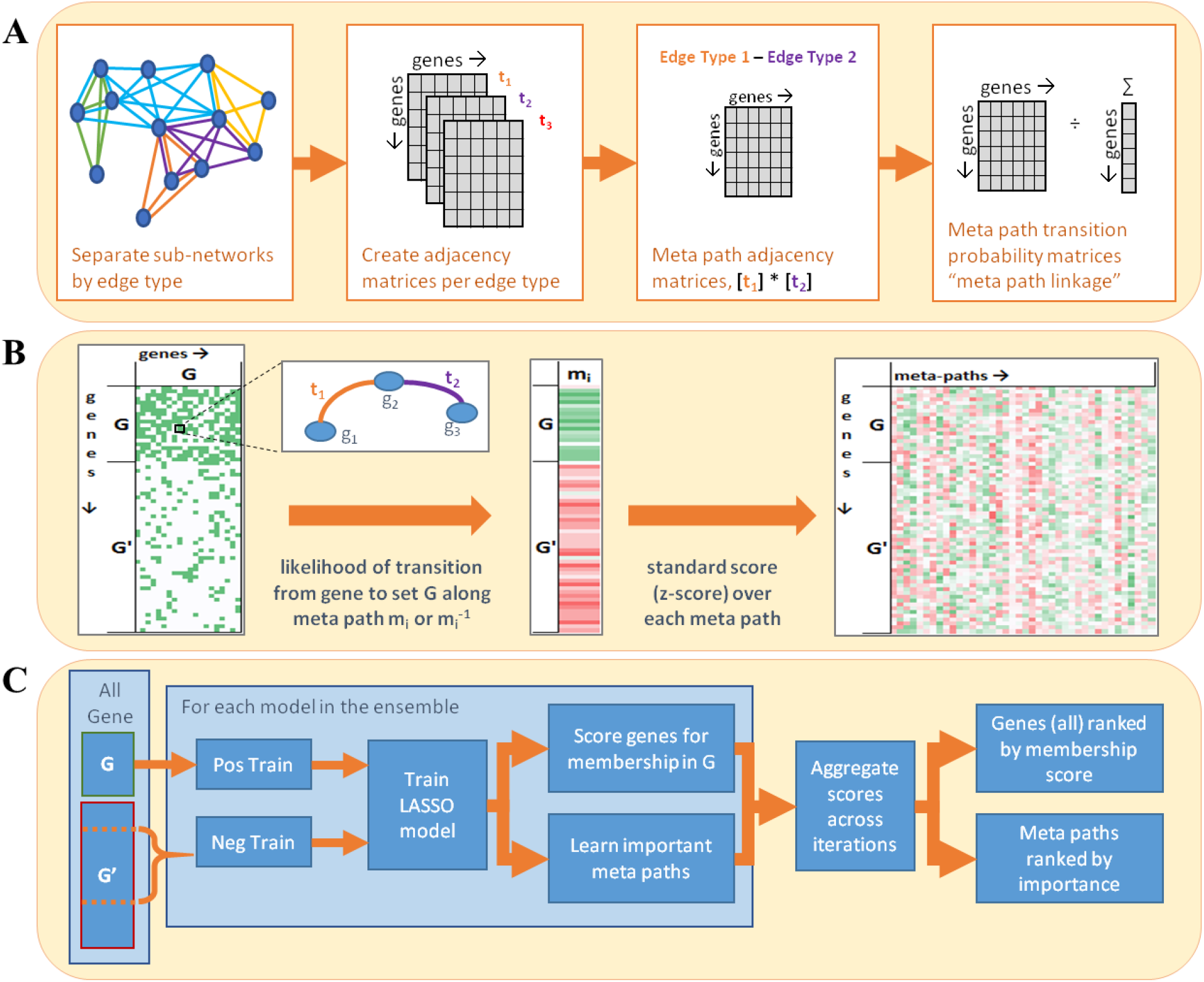
GeneSet MAPR Overview. A) GeneSet MAPR begins with a preprocessing step, in which, the weighted sub-networks of the heterogeneous network are converted into transition matrices for each meta path that describe the probability of connecting between two genes by following a meta path instance of the given type. B) When a query gene set is supplied, the linkage score is calculated between each gene and the query gene set for each type of meta path. C) Finally, these features are weighted in sparse regression models whose predictions are combined to create an aggregate score/rank for every gene relating it to the query gene set.

### Evaluations of Geneset MAPR

In order to evaluate MAPR’s effectiveness for the GSMP task, it was applied to 179 public gene sets selected from curated collections in the Molecular Signatures Database (MSigDB) (20), database of Genotypes and Phenotypes (dbGAP) (21), and Project Achilles (22) (**Supplementary Table 1A, Supplementary Note 1**). The MSigDB gene sets are constructed from the up- and down-regulated genes in 53 cancer studies conducted by multiple laboratories. Fifty-one gene sets selected from dbGaP represent genes associated with observable traits, from genome wide association studies. The 75 Achilles gene sets each correspond to a single cell line and contain the genes whose genetic knockout impacts the overall fitness of that cell line. While most of the results reported below pertain to these three collections of gene sets, we also performed similar evaluations on 320 additional gene sets from 8 other sources (**Supplementary Table 1B**) (23). As an evaluation metric, we used the Area Under the Receiver Operating Characteristic Curve (AUC) statistic associated with a four-fold cross-validation process where in each ‘fold’ 75% of the genes in a set were used for training and the ranks of the remaining 25% (hidden) genes were examined. MAPR performs strongly on all three collections – MSigDB, Achilles, and dbGaP and with an average AUC for the ranking of the left-out genes of 0.758, 0.714, and 0.677 respectively (Table 2A, **Supplementary Table 2A**). 79% (42 of 53) of the cancer gene sets from MSigDB and 40% (30 of 75) of the cell line gene sets from Achilles were especially well ranked (AUC > 0.7) (Table 2B). To further assess these performance metrics, we undertook systematic comparisons with three other methods, as described next.

**Table 2.**
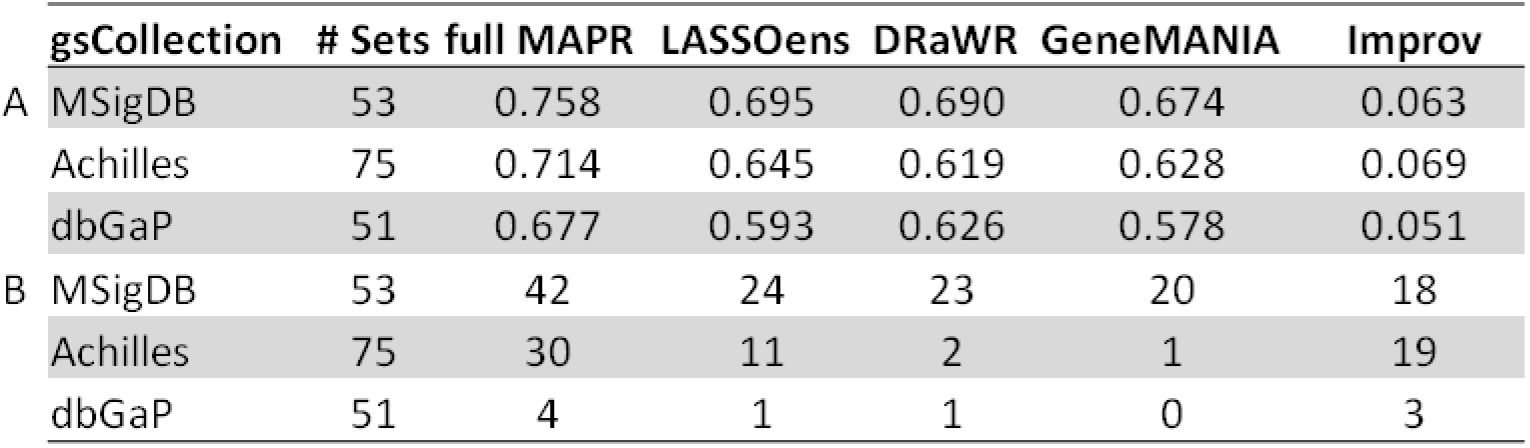
Average AUC Comparison by Method. The average AUC from four-fold cross validation is calculated for every gene set for four different methods (columns) using the same prior knowledge data: full MAPR, DRaWR, GeneMANIA, and LASSO Ensemble (similar to MAPR without meta paths). The analysis was performed on three separate gene set collections (rows) and the improvement of full MAPR over any of these three alternatives is reported in the ‘Improv’ column. A) reports the AUC values averaged across gene sets, B) reports the number of gene sets with AUC values greater than 0.7).

We first compared the performance of MAPR to a simplified version of itself, called ‘LASSO ensemble’, that adopts MAPR’s approach of aggregating predictions from an ensemble of LASSO regression models, but constructs the feature vectors describing genes differently. In particular, it describes each gene by a sparse vector whose features represent each unique annotation term or interactions of that gene. Importantly, unlike MAPR, it does not compute meta paths and ignores non-trivial connectivity patterns in the above-mentioned heterogeneous network, although it uses the same compendia of prior knowledge as MAPR. MAPR significantly outperforms the LASSO ensemble baseline, whose average AUC for MSigDB, Achilles, and dbGaP collections is lower than that of MAPR by 0.063, 0.069, and 0.084 respectively (Table 2A, Figure 3A, **Supplementary Table 2A**), demonstrating the advantage of meta path-specific linkage scores used by MAPR.

**Figure 3.**
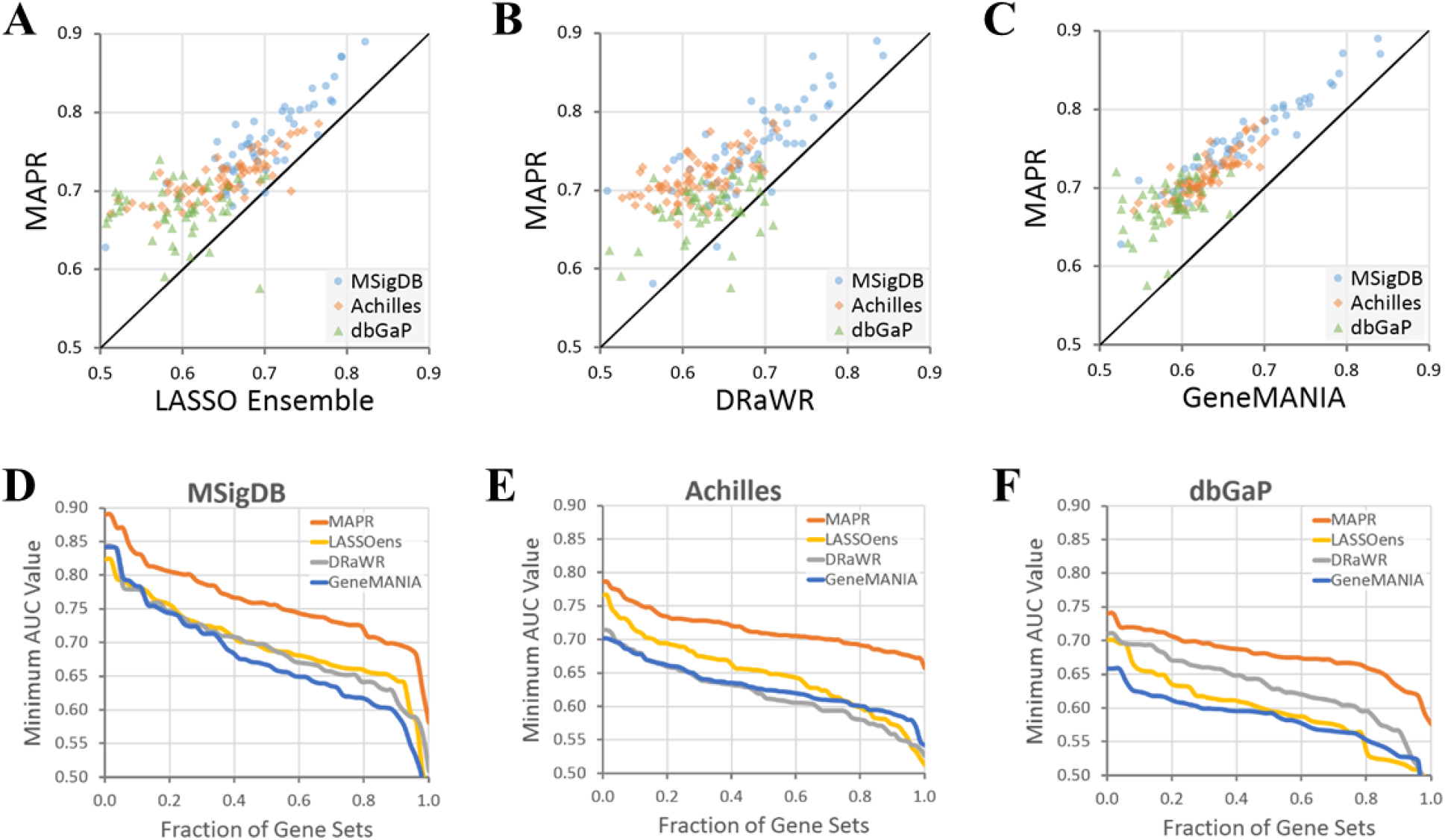
Comparison between MAPR and Other Methods. Scatterplots are drawn comparing the average cross-validation AUC for each gene set (plotted point) for our three selected collections (shape and color) of GeneSet MAPR to one of three alternative methods using the same heterogeneous network: A) LASSO ensemble, B) DRaWR, and C) GeneMANIA. Plots showing for each different method (colored lines), the percentage of gene sets (x-axis) achieving an average AUC or higher (y-axis) in one of our collections: D) MSigDB, E) Achilles, and F) dbGaP.

We next compared MAPR to another network-based method, called Discriminative Random Walk with Restart (DRaWR) (3), after adapting it for the GSMP task. DRaWR uses paths in the above-mentioned heterogeneous network to define similarity of individual genes to the query gene set, but it does so without distinguishing between edge types, treating the network as homogeneous after appropriate normalization. DRaWR also lacks the supervised learning aspect of MAPR’s algorithm, ranking genes based on their overall linkage to the query set rather than combining many meta path-specific linkage values via a trained model. We noted MAPR as achieving a higher four-fold cross-validation AUC on average than DRaWR across all tested gene set collections (Table 2A, **Supplementary Table 2A**). 96% of gene sets in MSigDB, 90% of sets in dbGaP, and 100% of sets of Achilles have an improved AUC with MAPR than with DRaWR (Figure 3B, **Supplementary Table 3**).

Our final comparisons were with the popular tool called GeneMANIA (6), which is capable of learning from heterogeneous networks in order to solve the GSMP problem, but differs from MAPR in its algorithm (see **Supplementary Note 2**). GeneMANIA ‘collapses’ the given heterogeneous network into a homogeneous one, whose edge weights are defined by a linear combination of the edge weights of the original networks. It then examines gene connectivity patterns in the resulting homogeneous network. In our runs of GeneMANIA using the same heterogeneous network as for MAPR (above), we found a clear difference in performance, with MAPR exhibiting clearly improved AUC over GeneMANIA for all tested collections (Table 2A, **Supplementary Table 2**, Figure 3C). These comparisons with DRaWR and GeneMANIA demonstrate the value of accounting for the heterogeneous nature of the network through use of meta paths. Viewing the above comparisons in another way, we noted that MAPR performs well (AUC > 0.7) on 18 additional genes sets (out of 53) in the MSigDB collection, 19 additional sets (of 75) in Achilles, and 3 additional sets (of 51) in the dbGaP collection than the second-best performing method for those collections (Figure 2D-F, Table 2B).

### Dissection of MAPR methodology

We investigated the contributions of major aspects of the MAPR methodology – choice of networks, meta paths, etc. – to its performance. The use of a heterogeneous network that combines prior knowledge from 11 different compendia was found to perform substantially better than when using any one of those compendia alone, represented as a homogeneous network (**Supplementary Table 4**). Individual compendia that led to the strongest performance include GO Biological Process, GO Cellular Component and STRING Text Mining. We also systematically assessed the impact of path lengths used in computing linkage scores, and noted that use of meta paths of length 2 and 3, i.e., going beyond direct connections (length 1), indeed improves the predictive performance of MAPR (**Supplementary Table 2, Supplementary Figure 1**).

We then examined the most informative meta paths (**Supplementary Table 5**) used by MAPR in the above evaluations and found that informative meta paths for MSigDB and dbGAP gene sets had a greater tendency to use ‘STRING text mining’ edges (a gene pair has this relationship if it co-occurs frequently in published abstracts) compared to meta paths useful for Achilles gene sets. In fact, the single most useful meta path for the MSigDB gene sets was ‘STRING_textmining – STRING_textmining’, indicating that gene pairs in these sets are most inter-related by way of their co-occurrence with a third gene in the literature. Since such co-occurrence may be due to a variety of biological or biochemical relationships, including those not covered by the compendia used here, this points to a variegated set of relationships connecting genes within a gene set. On the other hand, Gene Ontology relationships were commonly utilized by MAPR across all three collections, in agreement with the popular practice of reporting GO enrichments for differentially expressed (query) gene sets.

### Characteristics of predictable gene sets

We noticed that gene sets within the same collection are predictable to different degrees, e.g., AUC values of MAPR on gene sets in the MSigDB collection range between 0.58 (‘ACEVEDO LIVER CANCER WITH H3K9ME3’) and 0.89 (‘WOO LIVER CANCER RECURRENCE’) despite having similar origins/definitions, i.e., dysregulated genes from cancer studies. We thus asked if there are statistical properties shared by gene sets for which the GSMP task was more (or less) successful. We first noted that small gene sets (< 100 genes) are clearly less amenable to the GSMP task (**Supplementary Table 6**) by MAPR, as might be expected from the supervised nature of the algorithm. This is also partly the reason why gene sets from the dbGAP collection yielded a lower average AUC (above, Table 2A), since nearly half of its sets are in the 50-100 size range, while the MSigDB and Achilles collections have almost no gene set in this size category. Secondly, we noted that the gene sets more amenable to GSMP exhibited greater node degrees (counts of edges involving those genes) in the heterogeneous network. In particular, sets whose member genes had median degree more than 100 yielded significantly greater AUC values (by MAPR) than those with median degree less than 100 (**Supplementary Table 7**), with a Wilcoxon Rank Sum test p-value of 5.0E-35. This is expected, to an extent, since the MAPR method exploits network connections in order to solve the GSMP problem. At the same time, this does not mean that any gene set with high node degree will be amenable to the GSMP task; the predictor must still learn the right connectivity patterns that distinguish that gene set from all other genes. Finally, we repeated the above type of examination comprehensively by quantifying various properties of a gene set and computing the correlation between MAPR AUC value of that gene set and each of those properties (Table 3). In addition to node degree and set size, a higher proportion of genes related to cancer or potential drug targets appeared to indicate that a gene set is amenable to GSMP, presumably because these categories of genes tend to be better studied and hence connectivity patterns among them are more discernible.

**Table 3.**
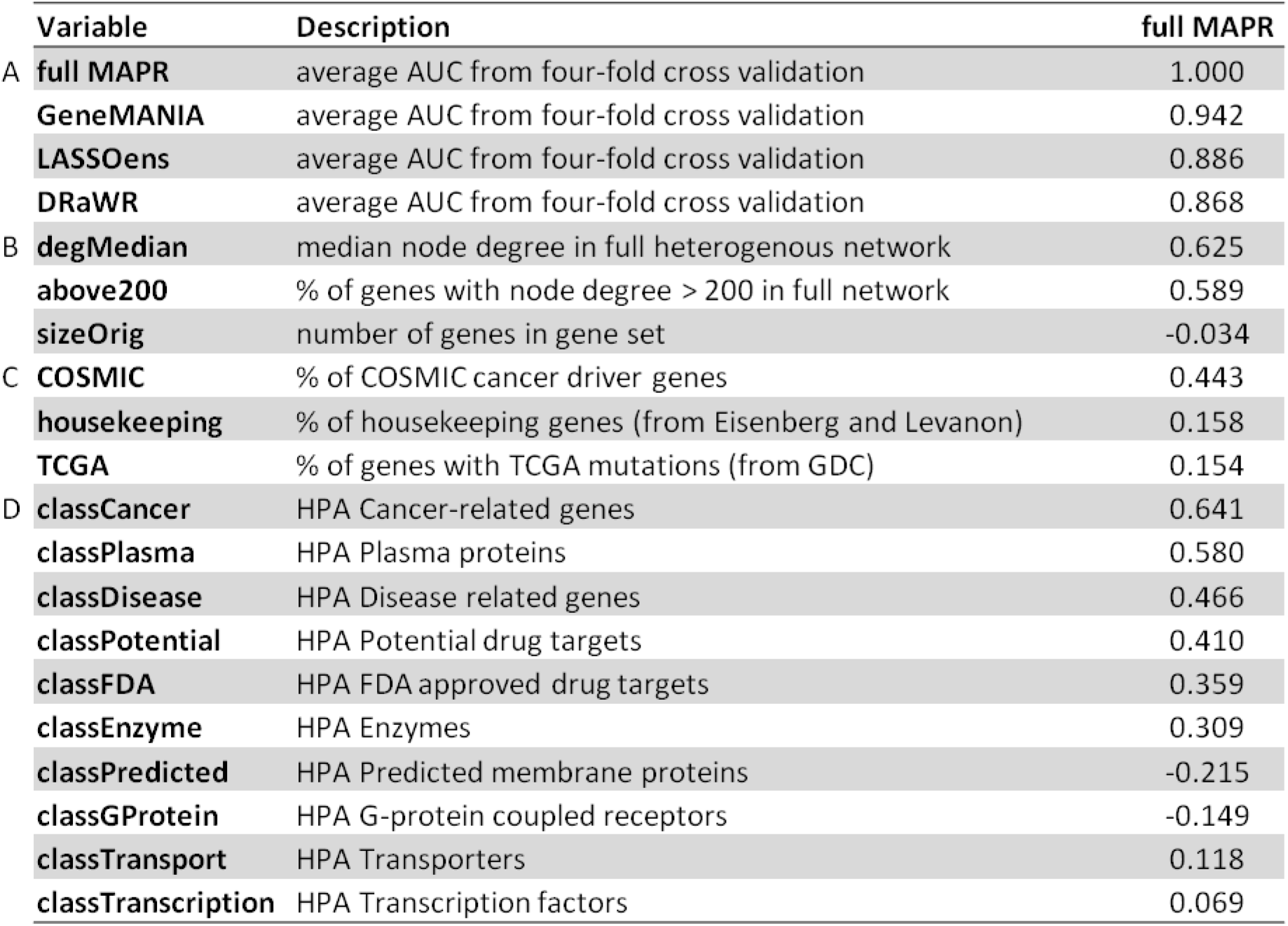
Relation between Gene Set Characteristics and MAPR Performance. The average AUC from four-fold cross validation is recorded for four competing methods (columns) for the 499 gene sets in this study. The Pearson correlation is calculated between these values and many different gene set characteristics (rows). This table show the correlation of performance A) between the methods, B) with network properties of the gene set, and with annotation percentages from C) specific databases (40,41) and D) the Human Protein Atlas (42).

### MAPR analysis of genes associated with patient response to chemotherapy: a case study

To illustrate the utility of GeneSet MAPR, we report a detailed analysis of a gene set that is differentially expressed between triple negative breast cancer patients who exhibited pathological complete response (pCR) to a chemotherapy regimen and those who did not (non-pCR) (24). This gene set was derived from a longitudinal study named ‘BEAUTY’ at the Mayo Clinic and we refer to it as ‘BEAUTY TRIPLE NEGATIVE RESPONSE’ or ‘BTNR’ gene set. As our query gene set, we selected 384 genes that had a differential expression p-value less than 0.05, were expressed (median raw genecounts > 32) and had at least two-fold change in between responders and non-responders. Of these genes, 323 are represented in the MAPR network.

We first noted that membership in the BTNR gene set is learnable by MAPR, as the four-fold cross validation AUC for the GSMP task on this set was 0.743 (**Supplementary Table 8**), with several meta paths shared across the trained models (**Supplementary Table 9**). MAPR was then trained on the entire set and used to rank genes for similarity to it. Of specific interest are genes ranked highly by MAPR, but which fell below the statistical cutoff values for differential expression that defined the BTNR set (**Supplementary Table 10**). Three families of genes showed a dramatically increased representation among MAPR’s top-ranked genes for the BTNR query set. While some members of these families were included in the BTNR set, they were not mentioned in the BEAUTY study (24). MAPR’s ranking brought them to attention. We briefly present our observations about these three families below. As a point of contrast, we also examined these families for their MAPR ranking when using other cancer gene sets (instead of the BTNR, see **Supplementary Note 3**) as query sets.

#### Claudins

This family is important to cell adhesion and flow of molecules in the intercellular space (25). While the BTNR gene set (384 genes) contained six claudins, MAPR trained on this gene set listed 20 claudins at a rank of 400 or better (**Supplementary Table 11A**). Three of these (claudin 5, claudin 23, claudin 19) were found to have SNVs in non-pCR patients, yet were omitted from the BTNR gene set constructed based on differential expression alone. Claudins were generally not ranked highly by MAPR when training on other cancer gene sets, suggesting a unique importance to the BTNR set. Indeed, claudin-low tumors have been identified as a distinct molecular subtype of breast cancer and are associated with poor prognosis (26,27).

#### Kallikreins

The BTNR gene set contained seven kallikreins, which are a type of serine proteases. MAPR listed those plus another seven in its top 400 list (**Supplementary Table 11B**). Five of the seven kallikrein genes from the BTNR set were also found by MAPR trained on other cancer gene sets. One of these, KLK2, is listed as a Tier 1 cancer gene by COSMIC but was not included in the BTNR gene set, because it is expressed at lower levels below the cutoff threshold. Kallikreins are cited by many studies as indicators of poor prognosis in cancer (28), and are often down-regulated and used as biomarkers in breast cancer patients (29). However, their presence as a family was not highlighted in the BEAUTY study (24), and MAPR’s gene ranking drew attention to their potential importance in this context.

#### Collagen Type N Alpha Chains

The BTNR gene set contained six collagens, and MAPR ranked these as well as an additional 31 in the top 400 (**Supplementary Table 11C**); most of these 37 had potentially significant levels of mutation among cancer patients according to TCGA. Of these, three (COL17A1, COL14A1, COL22A1) have a significant SNV (>7%) and/or CNV (>18%) rates between pCR and non-pCR patients, but were not included in the BTNR gene set. COL3A1, at rank 378 in the MAPR output, is listed as a Tier 2 gene by COSMIC, but was also not included in the BTNR set. Several collagen alpha-chains have been linked to cancer progression. Collagen Type VI (5 genes reported in the MAPR top 400) promotes both tumor progression and chemotherapy resistance, suggesting a feasible anticancer strategy wherein collagen VI is suppressed (30). Type V (3 genes in the MAPR top 400) alpha chains are higher in the desmoplasia surrounding tumors in breast tissue (31) and evidence in mice indicates Type V chains as potential targets for inhibiting tumor growth (32). (See **Supplementary Note 4** for more details.)

In summary, when presented with the ~400 gene query set of BTNR genes, MAPR re-ranks all genes based on connectivity patterns observed in the BTNR set, and the top 400 genes in this new ranking have a greatly enhanced presence of three gene families – claudins, kallikreins, and collagens – that are known for their association with poor prognosis. This observation motivated us to test more systematically for properties, e.g., GO terms, KEGG pathways and PFAM protein domains, enriched in the top ranked genes reported by MAPR trained on the BTNR set. We did this using the ‘Gene Set Enrichment Analysis’ (GSEA) (2) tool that can report if one gene set (the ‘property’, such as GO term) is statistically associated with a user-provided ranking of all genes or a user-provided gene set. Our proposed task is a variant of the standard GSEA application, where instead of testing enrichments in the BTNR gene set itself we sought enrichments in a MAPR-provided gene ranking based on similarity to the BTNR set. In a separate analysis (**Supplementary Table 12**), we tested this idea systematically on a handful of gene sets from our evaluation collection and found that the new ‘MAPR-GSEA’ method identifies meaningful properties of gene sets that are not necessarily found by the standard GSEA application.

On applying MAPR-GSEA to the BTNR gene set, the most strongly associated properties (by the Normalized Enrichment Score of GSEA) were found related to membrane/extracellular matrix, ion transport, cell adhesion, and signaling (Figure 4, **Supplementary Figure 2, Supplementary Table 13**). We asked if any of these identified properties were not ranked highly in a GSEA analysis of nine other related cancer gene sets (**Supplementary Table 14**). Indeed, properties such as protein digestion/absorption, the extracellular matrix, or membranes, which are related to the major gene families reported above, as well as properties related to signaling, ion transport, or cell adhesion, were found to be relatively unique in their association with the genes ranked highly by MAPR for similarity to the BTNR gene set.

**Figure 4.**
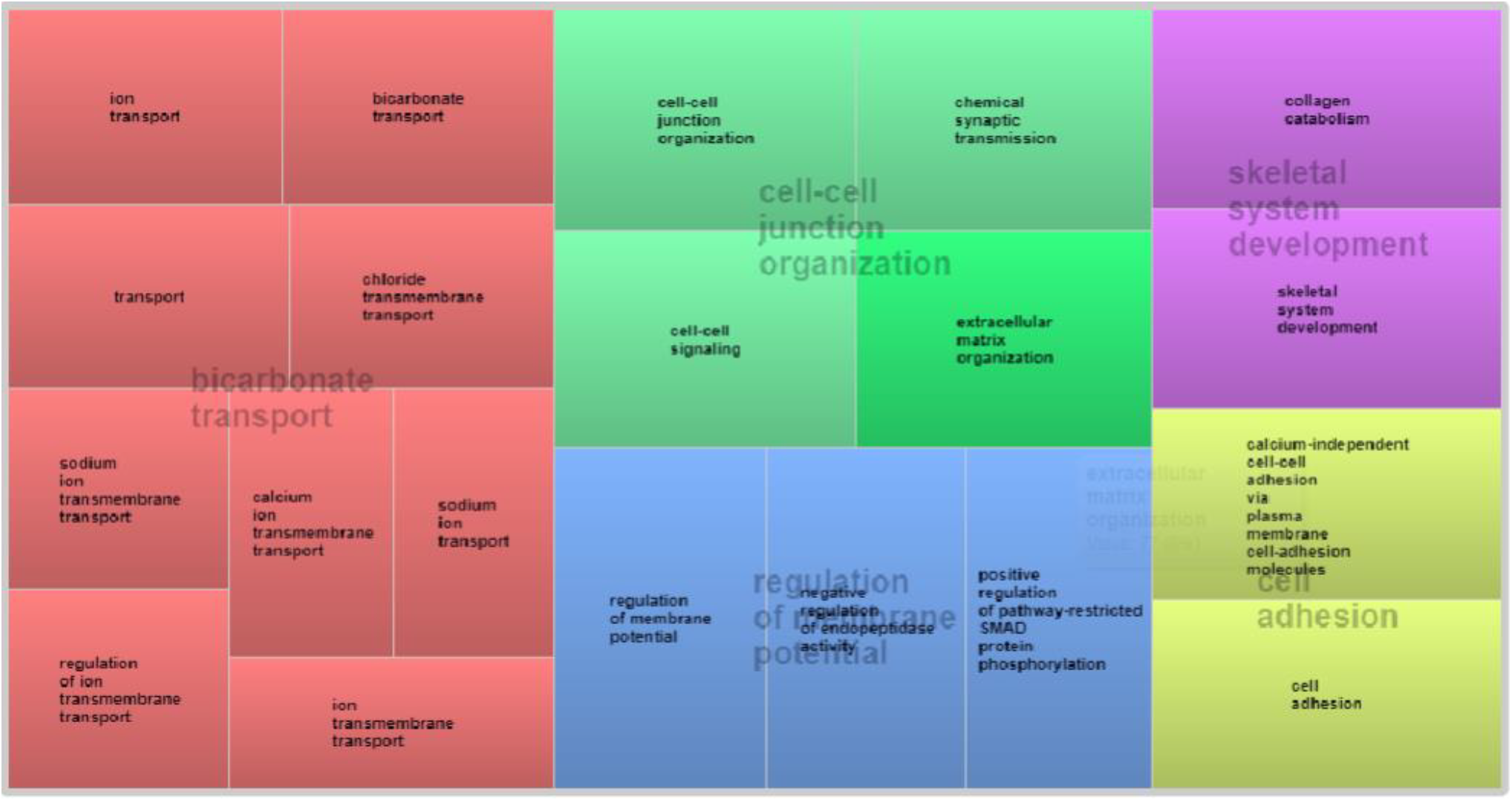
REViGO Visualization of BTNR Annotations. After producing a ranked gene list from MAPR for the BTNR gene set, we performed GSEA on that ranked list for related Gene Ontology terms. The REViGO (39) TreeMap visualization displayed above is for the top 50 GO annotations showing several functional clusters relating to ion (transmembrane) transport, signaling & extracellular matrix, cell adhesion, collagen, and membrane regulation.

## DISCUSSION

Gene set membership prediction has been the subject of significant research activity in the context of gene function prediction (4), where genes with a common Gene Ontology function annotation are used to identify other genes of the same function. Here, we study the GSMP problem in a far broader scope and investigate its feasibility on gene sets from a diverse array of sources, with the ultimate goal of allowing researchers to analyze any gene set of interest. We develop Geneset MAPR, a meta path-based algorithm for GSMP, and assess its performance against a popular current method called GeneMania, a network random walk-based method called DRaWR, as well as other suitable baselines. We investigate what kinds of prior knowledge are most useful for MAPR and what makes a gene set more amenable to the GSMP task. We also illustrate the utility of MAPR on a real gene set related to poor drug response in breast cancer patients, using this bioinformatics primitive to identify gene families and other annotations closely related to the gene set.

We believe that gene set membership prediction (GSMP) can be a powerful bioinformatics primitive with diverse uses in the analysis of gene sets. In addition to its above-mentioned uses, the accuracy of GSMP on a given gene set may be treated as a quantitative measure of the ‘modularity’ of the set, a concept that has been readily adopted in systems biology. Techniques for discovery of gene modules, for example from co-expression networks, often seek external confirmations or evaluations of the quality of those modules. In a recent DREAM challenge (33), a comprehensive assessment of module identification methods was attempted across a range of protein and gene networks. The challenge was to predict functional modularity over single and multiple networks. The evaluation relied on enrichment of predicted modules for genetic loci of diseases from GWAS studies. However, there can be significant commonality in tightly knit gene modules beyond their co-involvement in diseases. We note that a GSMP method such as MAPR may be used to evaluate modules, since one expects that a true gene module, comprising closely inter-related genes, will be amenable to the GSMP task. Since MAPR is able to utilize information from heterogeneous networks for this task, a simple cross-validation of MAPR on a candidate module can provide a richly informed test of its modularity, going beyond disease involvement.

Genome scale networks that capture high throughput experimental data (14), computationally discovered gene relationships (13,15,16,20) or manually curated gene properties (17,18,34) have become integral to knowledge encapsulation. This has enabled novel approaches to analyze and increase functional knowledge of genes and proteins under the implicit assumption that proximity between genes in the network is an indicator of their functional similarity – the ‘guilt-by-association’ principle (35). Gene networks can be reconstructed from a variety of sources, leading to a heterogeneous network that encapsulates some or all of these individual networks. The principled handling of heterogeneous information networks, however, remains an important challenge and the concept of gene-gene connectivity in such networks is ill defined. Geneset MAPR marks an attempt to address this major challenge in genomics, in the context of the GSMP task.

Furthermore, we believe that the concept of meta paths is powerful for heterogeneous network analysis, even outside of the genomics context. In many real world networks and graphs, the occurrence of sets of nodes in densely linked clusters, or communities, is a natural organizing principle. Whether this is as societal organization as groups in social networks, common hobby enthusiasts in a collaborative network or topically related pages in the World Wide Web, nodes serving similar functions or roles tend to form communities or modules in graphs. Detecting such communities from an unlabelled graph has been a subject of intense research in recent times. Given the widespread use of large graphs for data representation, and the emphasis on generating knowledge and insight from the graphs, community detection has become a seminal problem in network science today. Thus, our novel meta path-based approach to defining node relationships in heterogeneous networks will be of great relevance to the broader research community of network science.

## MATERIAL AND METHODS

### Basic definitions

A network is defined as nodes (genes) connected by edges (pairwise relationships or shared annotations of genes). In a homogeneous network, every edge is of the same type, specifying the same type of relationship, while a heterogeneous network contains more than one edge type. A path is a contiguous sequence of edges, where each successive pair of edges shares a node, that starts and ends at specified nodes. In a heterogeneous network, a meta path is an ordered sequence of edge types that matches the sequence of edge types of one or more individual paths in the network (12); such individual paths are called ‘instances’ of the meta path (Figure 1C,D).

Given a gene set, the goal of GeneSet MAPR is to score and rank genes by how related they are to the set. MAPR’s score is based on a collection of curated gene-gene relationships available *a priori*. Each type of relationship is recorded as a homogeneous network using genes as nodes and gene-gene relationships as edges. A heterogeneous network can then be defined as the union or ‘overlay’ of multiple homogeneous networks. In Results, we outlined how MAPR uses the heterogeneous network to define a ‘linkage’ between a gene and the given gene set, and how this vector is then used as a feature vector by a classifier, whose numeric output is the final score of the gene’s similarity to the given gene set. Here, we provide details of the steps mentioned in that outline with notation defined in (**Supplementary Figure 3**).

### Network preprocessing

In this step (Figure 2A), the heterogeneous network is pre-processed to create a ‘transition probability matrix’ corresponding to each meta path, for use in the ‘linkage’ score definition below. First, each homogeneous network is described by an adjacency matrix. An adjacency matrix is traditionally defined as a square matrix where (*i, j*)^th^ entry is 1 if there is an edge from node *i* to node j, and 0 otherwise. MAPR modifies this definition to account for networks where two genes may have multiple edges between them, and edges may have weights indicating the strength or confidence of the underlying relationship. Additionally, each edge is treated as undirected, regardless of whether the original relationship was symmetric or not. The adjacency matrix *A*^*t*^ for homogeneous network of edge type *t* is defined as:

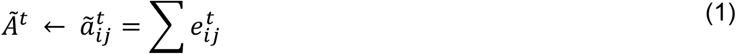

where 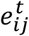 is all edge weights of type *t* that connect nodes *i* and *j*. Note that the adjacency values are normalized to a [0,1] range; this is done in order to give equal emphasis to each homogeneous network when defining linkage scores (below).

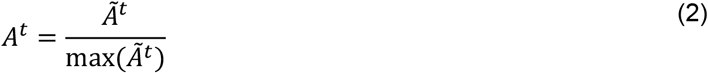

Next, the adjacency matrices for all edge types in a meta path *m* are utilized to construct a transition probability matrix corresponding to that meta path. Typically, a transition probability matrix parameterizes a random walk on a network: if one considers a random walk starting at a given node and taking a single step at random along any of its incident edges, then the (*i,j*)^th^ cell of the transition probability matrix prescribes the probability of transitioning from node *i* to *j* in one step of the walk (36). MAPR defines a transition probability matrix that captures connections between pairs of nodes, not as individual edges but as instances of a meta path. For each meta path *m*, it first defines an adjacency matrix 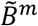 by multiplying adjacency matrices *A*^*t*^ in the order of edge types *t* specified by the meta path *m* [Eqn. 3 below]. Diagonal entries are set to 0 so as not to consider loops. Based on the new adjacency matrix, which represents a new graph where edges represent instances of meta path *m*, MAPR then computes a meta path transition matrix *B*^*m*^ [Eqn. 4 below]. Note that the meta path transition matrix is asymmetric: its (*i,j*)^th^ cell indicates the probability of transitioning from node (gene) *i* to *j* along meta path *m*, where *m* specifies an order of edge types.

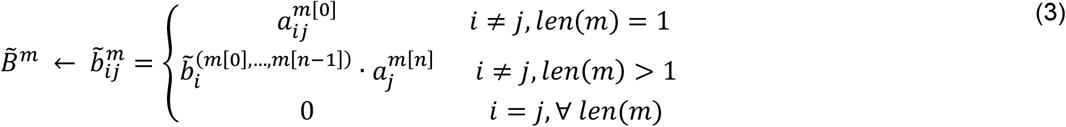

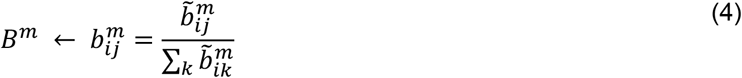

### ‘Linkage’ using meta paths

MAPR defines the linkage between two genes *i* and *j*, for a meta path *m*, as the transition probability from *i* to *j* along *m* or its inverse *m*^*−1*^, which is the sequence of edge types in opposite order as *m.* (As the edges in the network are treated as undirected, MAPR considers meta path connections in either direction to be equally important.) Then (Figure 2B), MAPR defines the linkage between gene *i* and gene set *G* as the probability of transitioning from *i* to any gene in *G*, along *m* or *m*^*−1*^ [Eqn. 5 below]. This is computed by considering the union of instances of meta path *m* between *i* and *j* or meta path *m*^*−1*^ between *i* and *j* (the latter being identical to instances of meta path *m* between *j* and *i*). The standardized linkage score, also known as the z-score (37), is defined by computing the mean and standard deviation of the linkage scores (for meta path *m*) of all genes, and then normalizing the linkage score of a gene by subtracting the mean and dividing by the standard deviation [Eqn. 6 below]. The standardized connected scores of a gene for all meta paths are the entries of a row vector that describes the gene’s relationship to the input gene set (7); row vectors of all genes are collected into a feature matrix *X* [Eqn. 7 below].

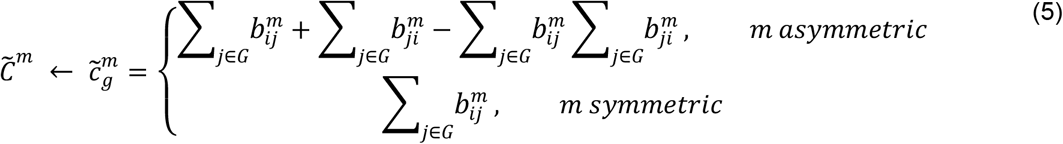

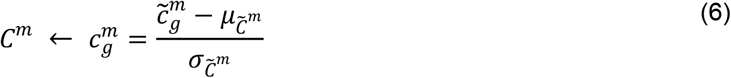

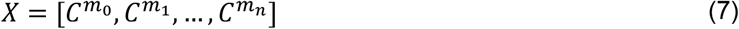

### LASSO regression for gene set membership prediction

MAPR uses the feature vectors (rows of *X*) of genes in the given gene set *G* (positive examples) and a sized-matched random subset of genes outside the set (negative samples) to train an ensemble of regression models that predict membership in *G* (Figure 2C). Each choice of the negative set defines a separate trained model. Training labels *y* are set to +1 for positive and 0 for negative samples. Each model uses LASSO regression (19), relying on L1-regularization to avoid over-fitting [Eqn. 8 below, *β*^{*i*}^ denotes the vector of learned coefficients of the *i*^*th*^ model]. The model performance score indicates how well the model labels the training data. For this, MAPR uses the general coefficient of determination (38), or r^2^ [Eqn. 9 below]. A score of 1 indicates every item was labelled correctly, 0 indicates every item was assigned the mean value, and a negative score indicates the model failed. A score of 0 or less results in attempting a new model based on a random sub-sample of the positive training set and a new negative set. In practice, this step has occurred rarely.

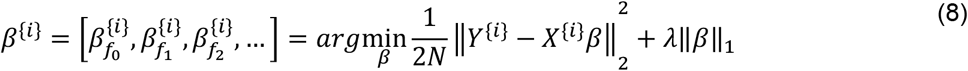

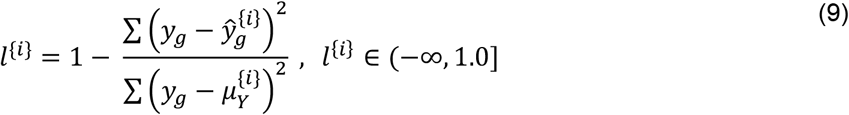

### Aggregating results from multiple models

Once a model has been trained, it is applied to predict a gene set membership score 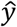 for every gene [Eqn. 10 below]. The score from each model is standardized, and then a weighted mean of the gene’s score from all models is computed; each model’s performance score is used as its weight in this step. [Eqn. 11]. This score, *s*_*g*_, is the reported similarity score of gene *g* for the given gene set. Similarly, an overall feature importance score is aggregated from the model coefficients [Eqn. 12].

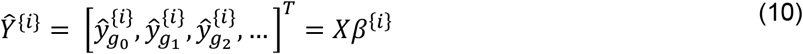

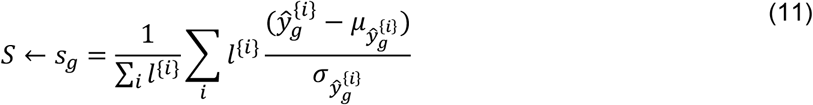

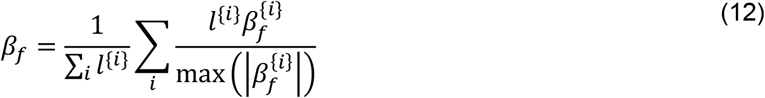

Thus, GeneSet MAPR reports two ranked lists related to the input gene set *G*: (i) a list of genes ordered by the similarity score *s*_*g*_, and (ii) a list of meta paths ordered by *β*_*f*_, which represents how important the meta path *f* was to the similarity score calculation.

### Evaluations and Comparisons

In each fold, 25% of the genes in the set were hidden and the remaining 75% were used as the query gene set for training MAPR. The trained predictor produced a ranked list of all genes and the position of the hidden genes in that ranking was used to calculate the AUC for that fold. The final AUC for each gene set is the average across the four folds.

For evaluations with DRaWR, we set the required ‘restart probability’ parameter to 0.3.

## AVAILABILITY

GeneSet MAPR is available as an open source software initiative available in the GitHub repository: https://github.com/KnowEnG/MetaPath_Analysis

## REFERENCES

1. Huang da, W., Sherman, B.T. and Lempicki, R.A. (2009) Systematic and integrative analysis of large gene lists using DAVID bioinformatics resources. Nat Protoc, 4, 44–57.

2. Subramanian, A., Tamayo, P., Mootha, V.K., Mukherjee, S., Ebert, B.L., Gillette, M.A., Paulovich, A., Pomeroy, S.L., Golub, T.R., Lander, E.S. et al. (2005) Gene set enrichment analysis: a knowledge-based approach for interpreting genome-wide expression profiles. Proc Natl Acad Sci U S A, 102, 15545–15550.

3. Blatti, C. and Sinha, S. (2016) Characterizing gene sets using discriminative random walks with restart on heterogeneous biological networks. Bioinformatics, 32, 2167–2175.

4. Pena-Castillo, L., Tasan, M., Myers, C.L., Lee, H., Joshi, T., Zhang, C., Guan, Y., Leone, M., Pagnani, A., Kim, W.K. et al. (2008) A critical assessment of Mus musculus gene function prediction using integrated genomic evidence. Genome Biol, 9 Suppl 1, S2.

5. Wang, S., Cho, H., Zhai, C., Berger, B. and Peng, J. (2015) Exploiting ontology graph for predicting sparsely annotated gene function. Bioinformatics, 31, i357–364.

6. Warde-Farley, D., Donaldson, S.L., Comes, O., Zuberi, K., Badrawi, R., Chao, P., Franz, M., Grouios, C., Kazi, F., Lopes, C.T. et al. (2010) The GeneMANIA prediction server: biological network integration for gene prioritization and predicting gene function. Nucleic Acids Res, 38, W214–220.

7. Sun, Y. and Han, J. (2012) Mining heterogeneous information networks: principles and methodologies. Synthesis Lectures on Data Mining and Knowledge Discovery, 3, 1–159.

8. Shi, C., Li, Y., Zhang, J., Sun, Y. and Philip, S.Y. (2017) A survey of heterogeneous information network analysis. IEEE Transactions on Knowledge and Data Engineering, 29, 17–37.

9. Cornish, A.J. and Markowetz, F. (2014) SANTA: quantifying the functional content of molecular networks. PLoS Comput Biol, 10, e1003808.

10. Hou, J.P. and Ma, J. (2014) DawnRank: discovering personalized driver genes in cancer. Genome Med, 6, 56.

11. Hofree, M., Shen, J.P., Carter, H., Gross, A. and Ideker, T. (2013) Network-based stratification of tumor mutations. Nat Methods, 10, 1108–1115.

12. Sun, Y., Han, J., Yan, X., Yu, P.S. and Wu, T. (2011) Pathsim: Meta path-based top-k similarity search in heterogeneous information networks. Proceedings of the VLDB Endowment, 4, 992–1003.

13. Szklarczyk, D., Franceschini, A., Wyder, S., Forslund, K., Heller, D., Huerta-Cepas, J., Simonovic, M., Roth, A., Santos, A., Tsafou, K.P. et al. (2015) STRING v10: protein-protein interaction networks, integrated over the tree of life. Nucleic Acids Res, 43, D447–452.

14. Chatr-Aryamontri, A., Oughtred, R., Boucher, L., Rust, J., Chang, C., Kolas, N.K., O’Donnell, L., Oster, S., Theesfeld, C., Sellam, A. et al. (2017) The BioGRID interaction database: 2017 update. Nucleic Acids Res, 45, D369–D379.

15. Finn, R.D., Coggill, P., Eberhardt, R.Y., Eddy, S.R., Mistry, J., Mitchell, A.L., Potter, S.C., Punta, M., Qureshi, M., Sangrador-Vegas, A. et al. (2016) The Pfam protein families database: towards a more sustainable future. Nucleic Acids Res, 44, D279–285.

16. Altschul, S.F., Gish, W., Miller, W., Myers, E.W. and Lipman, D.J. (1990) Basic local alignment search tool. J Mol Biol, 215, 403–410.

17. Ashburner, M., Ball, C.A., Blake, J.A., Botstein, D., Butler, H., Cherry, J.M., Davis, A.P., Dolinski, K., Dwight, S.S., Eppig, J.T. et al. (2000) Gene ontology: tool for the unification of biology. The Gene Ontology Consortium. Nat Genet, 25, 25–29.

18. Kanehisa, M., Furumichi, M., Tanabe, M., Sato, Y. and Morishima, K. (2017) KEGG: new perspectives on genomes, pathways, diseases and drugs. Nucleic Acids Res, 45, D353–D361.

19. Tibshirani, R. (1996) Regression shrinkage and selection via the lasso. Journal of the Royal Statistical Society. Series B (Methodological), 267–288.

20. Liberzon, A., Subramanian, A., Pinchback, R., Thorvaldsdottir, H., Tamayo, P. and Mesirov, J.P. (2011) Molecular signatures database (MSigDB) 3.0. Bioinformatics, 27, 1739–1740.

21. Tryka, K.A., Hao, L., Sturcke, A., Jin, Y., Wang, Z.Y., Ziyabari, L., Lee, M., Popova, N., Sharopova, N., Kimura, M. et al. (2014) NCBI’s Database of Genotypes and Phenotypes: dbGaP. Nucleic Acids Res, 42, D975–979.

22. Cowley, G.S., Weir, B.A., Vazquez, F., Tamayo, P., Scott, J.A., Rusin, S., East-Seletsky, A., Ali, L.D., Gerath, W.F., Pantel, S.E. et al. (2014) Parallel genome-scale loss of function screens in 216 cancer cell lines for the identification of context-specific genetic dependencies. Sci Data, 1, 140035.

23. Chen, E.Y., Tan, C.M., Kou, Y., Duan, Q., Wang, Z., Meirelles, G.V., Clark, N.R. and Ma’ayan, A. (2013) Enrichr: interactive and collaborative HTML5 gene list enrichment analysis tool. BMC Bioinformatics, 14, 128.

24. Goetz, M.P., Kalari, K.R., Suman, V.J., Moyer, A.M., Yu, J., Visscher, D.W., Dockter, T.J., Vedell, P.T., Sinnwell, J.P., Tang, X. et al. (2017) Tumor Sequencing and Patient-Derived Xenografts in the Neoadjuvant Treatment of Breast Cancer. J Natl Cancer Inst, 109.

25. Tabaries, S. and Siegel, P.M. (2017) The role of claudins in cancer metastasis. Oncogene, 36, 1176–1190.

26. Prat, A., Parker, J.S., Karginova, O., Fan, C., Livasy, C., Herschkowitz, J.I., He, X. and Perou, C.M. (2010) Phenotypic and molecular characterization of the claudin-low intrinsic subtype of breast cancer. Breast Cancer Res, 12, R68.

27. Dias, K., Dvorkin-Gheva, A., Hallett, R.M., Wu, Y., Hassell, J., Pond, G.R., Levine, M., Whelan, T. and Bane, A.L. (2017) Claudin-Low Breast Cancer; Clinical & Pathological Characteristics. PLoS One, 12, e0168669.

28. Borgono, C.A. and Diamandis, E.P. (2004) The emerging roles of human tissue kallikreins in cancer. Nat Rev Cancer, 4, 876–890.

29. Paliouras, M., Borgono, C. and Diamandis, E.P. (2007) Human tissue kallikreins: the cancer biomarker family. Cancer Lett, 249, 61–79.

30. Chen, P., Cescon, M. and Bonaldo, P. (2013) Collagen VI in cancer and its biological mechanisms. Trends Mol Med, 19, 410–417.

31. Barsky, S.H., Rao, C.N., Grotendorst, G.R. and Liotta, L.A. (1982) Increased content of Type V Collagen in desmoplasia of human breast carcinoma. Am J Pathol, 108, 276–283.

32. Huang, G., Ge, G., Izzi, V. and Greenspan, D.S. (2017) alpha3 Chains of type V collagen regulate breast tumour growth via glypican-1. Nat Commun, 8, 14351.

33. Choobdar, S., Ahsen, M.E., Crawford, J., Tomasoni, M., Lamparter, D., Lin, J., Hescott, B., Hu, X., Mercer, J., Natoli, T. et al. (2018) Open Community Challenge Reveals Molecular Network Modules with Key Roles in Diseases. bioRxiv, 265553.

34. Fabregat, A., Jupe, S., Matthews, L., Sidiropoulos, K., Gillespie, M., Garapati, P., Haw, R., Jassal, B., Korninger, F., May, B. et al. (2018) The Reactome Pathway Knowledgebase. Nucleic Acids Res, 46, D649–D655.

35. Li, Z.-C., Huang, M.-H., Zhong, W.-Q., Liu, Z.-Q., Xie, Y., Dai, Z. and Zou, X.-Y. (2015) Identification of drug–target interaction from interactome network with ‘guilt-by-association’principle and topology features. Bioinformatics, 32, 1057–1064.

36. Gagniuc, P.A. (2017) Markov Chains: From Theory to Implementation and Experimentation. John Wiley & Sons.

37. Norman, G.R. and Streiner, D.L. (2008) Biostatistics: the bare essentials. PMPH-USA.

38. Devore, J.L. (2011) Probability and Statistics for Engineering and the Sciences. Cengage learning.

39. Supek, F., Bosnjak, M., Skunca, N. and Smuc, T. (2011) REVIGO summarizes and visualizes long lists of gene ontology terms. PLoS One, 6, e21800.

40. Futreal, P.A., Coin, L., Marshall, M., Down, T., Hubbard, T., Wooster, R., Rahman, N. and Stratton, M.R. (2004) A census of human cancer genes. Nat Rev Cancer, 4, 177–183.

41. Eisenberg, E. and Levanon, E.Y. (2013) Human housekeeping genes, revisited. Trends Genet, 29, 569–574.

42. Uhlen, M., Fagerberg, L., Hallstrom, B.M., Lindskog, C., Oksvold, P., Mardinoglu, A., Sivertsson, A., Kampf, C., Sjostedt, E., Asplund, A. et al. (2015) Proteomics. Tissue-based map of the human proteome. Science, 347, 1260419.

