## Supplementary Notes, Figures, and Table Legends for "Gene Sets Analysis using Network Patterns"

### Supplementary Note 1. Creation of Gene Set Collections

Our analysis of MAPR began with our three primary gene set collections: MSigDB, dbGaP, and Achilles. For MSigDB, we selected all of their cancer-related differentially expressed gene sets that had up and down regulated gene sets with a minimum of 50 genes. In total, there were 53 of these gene sets that combined the up and down regulated genes. We wanted to investigate two additional gene set collections that were very different from differential gene expression in tumors. DbGaP gene sets are mostly based on genomic studies among populations. Achilles gene sets are derived from gene knockdowns and cell line survivability. For dbGaP, we selected their 51 largest gene sets (all gene sets with size greater than 70). For Achilles, there were 216 sets with size >70, so we randomly sampled 75 of the gene sets for analysis.

For our secondary gene set collections, we had a simple procedure. For each additional gene set collection, we found all the gene sets that had at least 75 and no more than 2000 genes, and then randomly selected 40 gene sets from that collection.

### Supplementary Note 2. GeneMANIA Algorithm and Analysis

GeneMANIA (1) takes a user-provided list of genes and extends that list with genes that share properties or similar functions, displaying the weights of the networks used to define this similarity. The user has the option to limit which networks are used to find similar genes, as well as define the method used to weight the networks. GeneMANIA's networks are drawn from publicly available databases. Those not already available as gene-gene networks are converted into functional association networks. For longer gene lists, GeneMANIA defaults to a weighting method where it attempts to maximize the interactions between genes in the user list while minimizing the interaction of genes on the list with those that are not. For shorter gene lists where this is deemed impractical, by default GeneMANIA attempts to reproduce the Gene Ontology biological processes co-annotation patterns rather than the gene list. Genes not in the user list are assigned a score used to rank the non-query genes by relevance.

To compare methods, we imported our own networks into the GeneMANIA algorithm. We provided each edge type as its own edge list, just as they are provided to GeneSet MAPR prior to any processing. Thus, GeneMANIA used the same 11 subnetworks as GeneSet MAPR. We provided the same gene lists for GeneMANIA as were used by GeneSet MAPR; the cross-fold permutations used by MAPR were exported as text files and imported by GeneMANIA. The only default setting we overrode was the maximum number of related genes returned by the algorithm. We retuned all genes in the network, sorted by GeneMANIA's similarity score in order to produce AUC values.

### Supplementary Note 3. Nine Cancer-related Gene Sets

In order to assess whether the top ranking genes and Gene Ontology terms were specific to the BTNR gene set, we gathered nine additional cancer related gene sets to compare against. Seven of these gene sets were selected from the MSigDB (2) collection and related to these seven studies: "POOLA_INVASIVE_BREAST_CANCER", "SMID_BREAST_CANCER_BASAL", "SMID_BREAST_CANCER_LUMINAL_B", "SOTIRIOU_BREAST_CANCER_GRADE_1_VS_3", "ZHANG_BREAST_CANCER_PROGENITORS", "HUMMERICH_SKIN_CANCER_PROGRESSION", and

"WOO_LIVER_CANCER_RECURRENCE". The final two cancer related gene sets were created from TCGA gene expression data. We downloaded the gene expression and clinical information from the UCSC Cancer Genome Browser (3) for the breast invasive carcinoma (BRCA) project. We then calculated the significance of differentially expressed genes using a simple t-test in two ways: 1) between samples with the "Primary Tumor" vs "Metastatic" sample_type and 2) between samples with the "Primary Tumor" vs "Solid Tissue Normal". We then selected the 323 (size-matched to BTNR gene set) most differentially expressed genes in each of the two comparisons for our final cancer-related gene sets.

### Supplementary Note 4. Details on Collagen Type N Alpha Chains

Aberrations in Type I and Type III chains have been linked to malignant tumors, via the formation of readily-degradable collagen bundles (4). Both Type 1 genes were highly ranked, with COL1A2 (rank 267) showing a notable difference in mutation rates between pCR and non-pCR TNBC patients. COL3A1 (rank 378) showed a significant general rate of mutation and is listed as a Tier 2 gene by COSMIC. Types XIV (rank 155) and XXII (rank 59) showed significant rates of mutation among pCR TNBC patients, as opposed to non-pCR, although neither exhibited the required DE and p-value to be included in BTNR. Interestingly, Type XI alpha 1 (COL11A1) was ranked highly by MAPR (rank 170), while the remaining members of type XI were not. COL11A1 has been identified as an accurate marker for invasive breast carcinoma lesions (5), and as a potential target for general cancer therapy (6).

### Supplementary Figures

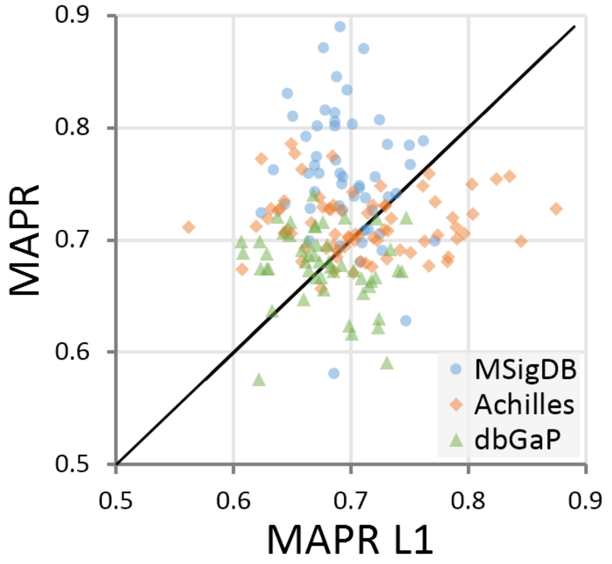

**Supplementary Figure 1. MAPR vs MAPR L1 Scatterplot**
Scatterplot comparing the average cross-validation AUC for each gene set (plotted point) for our three selected collections (shape and color) of GeneSet MAPR to a version of MAPR that only creates feature for meta paths of length one.

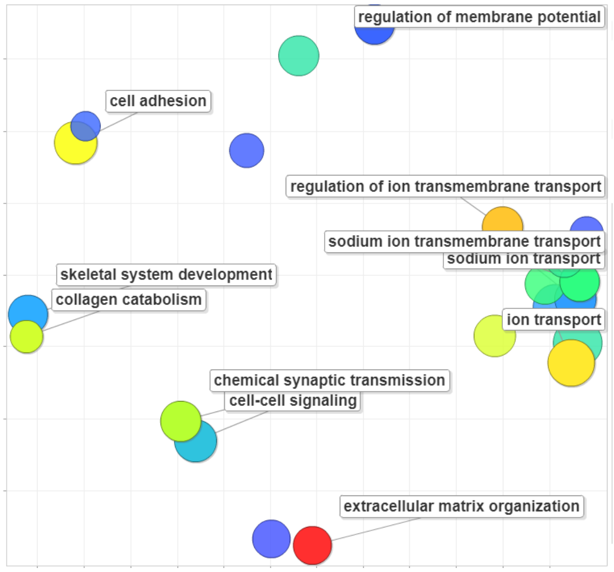

**Supplementary Figure 2. REViGO visualizations of BTNR Annotations.**
After producing a ranked gene list from MAPR for the BTNR gene set, we performed GSEA on that ranked list for related Gene Ontology terms. Gene Ontology terms clustered in REViGO's 2D semantic space, colored by GSEA enrichment score (red his higher, blue is lower). Transport terms are clustered to the right while signaling, collagen, extracellular matrix, and cell adhesion are in groups to the left.

| Symbol | Description | Symbol | Description |
|---|---|---|---|
| | | $e_i^t$ | Edge instance, of type t |
| $\boldsymbol{N^t}$ | Gene-gene network of type t | $t_i$ | Edge type i, from all available edge types |
| $\boldsymbol{I_{g \in G}}$ | Indicator that gene g is in gene set G | $g_i$ | Gene i |
| $\boldsymbol{M}$ | Collection of meta paths | $m_i$ | meta path i |
| $\boldsymbol{A^t}$ | Edge adjacency matrix of type t | $a_{ij}^t$ | Adjacency of gene i to gene j along edges of type t |
| $\boldsymbol{B^m}$ | Transition matrix for meta path of type m | $b_{ij}^m$ | Transition probability of gene i to gene j along meta path m |
| $\boldsymbol{C^m}$ | Linkage vector for meta path type m | $c_g^m$ | Linkage of gene g to set G along meta path m |
| $\boldsymbol{X}^{\{i\}}$ | Training Data / Gene Feature Vectors | $x_i^{\{+\}}, x_i^{\{-\}}$ | Pos, Neg training sets used for LASSO model i |
| $\boldsymbol{Y}^{\{i\}}$ | Training Labels / Gene Labels (in/out of set) | $y_g^{\{i\}}$ | Label of each gene used to train LASSO model i |
| $\boldsymbol{S}$ | Similarity score | $s_g$ | Standardized similarity score for gene g from model i |

**Supplementary Figure 3. List of MAPR Algorithm Symbols**
List of symbols and descriptions used to describe steps in the MAPR method.

### Supplementary Tables

**Supplementary Table 1. Details for Gene Set Collections.**
For each gene set collection, lists the alias and source descriptions, as well as the number of gene sets, and the average, minimum, and maximum size of those gene sets. Divided between A) three primary collections and B) eight additional collections.

**Supplementary Table 2. Average AUC for Gene Set Collections.**
Four-fold cross-validation was performed for each gene set and each possible method (column), and the area under the ROC curve (AUC) was calculated. The average of these gene set cross-validation AUC averages was calculated for each gene set collection (rows). Four different methods using the same prior knowledge data are compared (MAPR, DRaWR, GeneMANIA, and LASSOens (similar to MAPR without meta paths). Also, four additional methods were compared using MAPR with restricted meta paths (L1: paths of length one only, L2: paths of length two only, L3: paths of length three only, L1&2: only paths of length one or two). The analysis was done for the A) three primary gene set collections and B) eight additional gene set collections.

**Supplementary Table 3. Average AUCs for Gene Sets.**
Four-fold cross-validation was performed for each gene set and each possible method (column), and the area under the ROC curve (AUC) was averaged. Four different methods using the same prior knowledge data are compared (MAPR, DRaWR, GeneMANIA, and LASSOens (similar to MAPR without meta paths). Also, four additional methods were compared using MAPR with restricted meta paths (L1: paths of length one only, L2: paths of length two only, L3: paths of length three only, L1&2: only paths of length one or two). Finally, eleven additional methods using MAPR where compared using only one prior data source (GO-BP: Gene Ontology Biological Process, GO-CC: Gene Ontology Cellular Component, GO-MF: Gene Ontology Molecular Function, Homol: Blastp protein homology, KEGG: KEGG pathways, PFam: PFam protein domain annotations, PPI Direct: direct protein interactions, Gen Int: genetic interactions, PPI Physical: physical protein interactions, Co-Ex: STRING co-expression, Text: STRING Text Mining. There are also column for the gene set group, collection, size, and median degree in in the full MAPR network.

**Supplementary Table 4. Average AUC for Gene Set Collections by Homogenous Types.**
Four-fold cross-validation was performed for each gene set and each possible method (column), and the area under the ROC curve (AUC) was calculated. The average of these gene set cross-validation AUC averages was calculated for each gene set collection (rows). Besides the full, heterogeneous MAPR run with meta paths composed of all edge types (col E), eleven additional setups using MAPR with homogeneous meta paths using only one data source (GO-BP: Gene Ontology Biological Process, GO-CC: Gene Ontology Cellular Component, GO-MF: Gene Ontology Molecular Function, Homol: Blastp protein homology, KEGG: KEGG pathways, PFam: PFam protein domain annotations, PPI-Dir: direct protein interactions, Genetic: genetic interactions, PPI-Phys: physical protein interactions, Co-Ex: STRING co-expression, Text: STRING Text Mining) were evaluated. A) shows the three selected collections, B) the remaining gene set collections, and C) averages across all gene sets. D) For the selected gene set collections, we also show these statistics when the MAPR method was run with a heterogeneous network containing all but the edge type described by the column header.

**Supplementary Table 5. Top Meta paths Per Gene Set Collection.**
Each run of MAPR produces LASSO weights for combining the meta path features to predict gene set membership. These weights are min-max normalized and then averaged across folds and gene sets for

each meta path. For our three primary gene set collections, we report the top 10 meta paths by this score ('Avg Weight'), the length of the meta path ('MP Length'), and the percentage of runs in the gene set collection that learned a non-zero LASSO weight for this meta path ('Percent%').

**Supplementary Table 6. Average AUC by Gene Set Size.**
All gene sets from our 11 gene set collections were placed into one of six size bins based on the number of genes it contains. Each bin is described by its minimum "Gene Set Size". The average AUCs from four-fold cross-validation on each gene set are averaged for each size bin (row) and method (column).

**Supplementary Table 7. Average AUC by Gene Set Median Node Degree.**
All gene sets from our 11 gene set collections were placed into one of eight size bins based on the median degree in the heterogeneous MAPR network of its genes. Each bin is described by its minimum "Median Degree Size". The average AUCs from four-fold cross-validation on each gene set are averaged for each degree bin (row) and method (column).

**Supplementary Table 8. Average AUC for BTNR Gene Set by Homogenous Types.**
Shows the average of four-fold cross-validation area under the ROC curve (AUC) for the BTNR gene set using the full, heterogeneous MAPR run with meta paths composed of all edge types ('Combined') and eleven additional setups using MAPR with homogeneous meta paths using only one data source (GO-BP: Gene Ontology Biological Process, GO-CC: Gene Ontology Cellular Component, GO-MF: Gene Ontology Molecular Function, Homol: Blastp protein homology, KEGG: KEGG pathways, PFam: PFam protein domain annotations, PPI-Dir: direct protein interactions, Genetic: genetic interactions, PPI-Phys: physical protein interactions, Co-Ex: STRING co-expression, Text: STRING Text Mining).

**Supplementary Table 9. Top Meta paths for BTNR.**
Each run of MAPR produces LASSO weights for combining the meta path features to predict gene set membership. These weights are min-max normalized and then averaged across folds and gene sets for each meta path. For the BTNR gene set, we report the top 10 meta paths by this score ('Avg Weight'), the length of the meta path ('MP Length'), and the percentage of runs in the gene set collection that learned a non-zero LASSO weight for this meta path ('Percent%').

**Supplementary Table 10. Gene Ranking for BTNR Set.**
The ranking of genes ('MAPR Rank - BTNR') by the full version of MAPR with the combined heterogeneous network on the BEAUTY Triple Negative Response (BTNR) gene set. In MAPR Results, we show the MAPR rank and combined prediction score ('MAPR Score - BTNR') and an indicator for if the gene is in the top 400 results ('MAPR BTNR Top400'). We also show an indicator if the gene was in the original differential expression gene set ('Orig Set') as well as the differential expression p-value ('DE pval'). We examine other data from the BEAUTY study, the log fold change of the gene expression ('BTNR logFC'), the median gene expression level ('BTNR Median Expr'), the percentage of occurrences of nonsynonymous SNVs ('BTNR SNVs') and CNVs ('BTNR CNV') in the cohort. We also collected the percentage of samples in TCGA with a somatic mutation in each gene for breast cancer only ('SM across BRCA') and across all cancer types ('SM across PanCan'). We also check whether the genes are in COSMIC Cancer Gene Census ('CGC Tier'). Finally, we rerun MAPR on nine other cancer gene sets for MSigDB and find the number of times a gene occurs ('Num Sets') and its median rank ('Median MAPR Rank').

**Supplementary Table 11. Gene Families Returned by MAPR.**

Shows the gene evidences for three different families discovered by the MAPR ranking of genes on the BEAUTY Triple Negative Response (BTNR) gene set; A) claudins, B) kallikreins, and c) collagen type alpha chains. We show the MAPR rank ('MAPR Rank - BTNR') and combined prediction score ('MAPR Score - BTNR') and an indicator for if the gene is in the top 400 results ('MAPR BTNR Top400'). We also show an indicator if the gene was in the original differential expression gene set ('Orig Set') as well as the differential expression p-value ('DE pval'). We examine other data from the BEAUTY study, the log fold change of the gene expression ('BTNR logFC'), the median gene expression level ('BTNR Median Expr'), the percentage of occurrences of nonsynonymous SNVs ('BTNR SNVs') and CNVs ('BTNR CNV') in the cohort. We also collected the percentage of samples in TCGA with a somatic mutation in each gene for breast cancer only ('SM across BRCA') and across all cancer types ('SM across PanCan'). We also check whether the genes are in COSMIC Cancer Gene Census ('CGC Tier'). Finally, we rerun MAPR on nine other cancer gene sets for MSigDB and find the number of times a gene occurs ('Num Sets') and its median rank ('Median MAPR Rank').

**Supplementary Table 12. Annotations Returned by MAPR GSEA.**

Association tests with Gene Ontology terms were performed for seven gene sets in the MSigDB collection in one of two ways: with overlap enrichment using the one-sided Fisher exact test ,"Fisher", and for positive enrichment using GSEA on the full gene ranking produced by MAPR for the gene set, "MAPR GSEA". The table shows for each gene set, "Gene Set Name", the Gene Ontology terms ("GO Term") that were in the top 15 enriched terms using the MAPR GSEA method and not in the top 100 enriched terms using the Fisher method. Only GO terms of size, "SetSize", between 10 and 500 genes were considered. For MAPR GSEA, the term rank and normalized enrichment score (NES) are shown. For Fisher, the rank and the p-value are listed.

**Supplementary Table 13. Annotations Returned by MAPR GSEA for BTNR.**

Association tests with Gene Ontology terms were performed for the BTNR gene set in one of two ways: with overlap enrichment using the one-sided Fisher exact test ,"Fisher", and for positive enrichment using GSEA on the full gene ranking produced by MAPR for the gene set, "MAPR GSEA". The table shows the Gene Ontology terms ("GO Term") that were in the top 50 enriched terms using the MAPR GSEA method. Only GO terms of size ("SetSize") between 10 and 500 genes were considered. For MAPR GSEA, the term rank and normalized enrichment score (NES) are shown. For Fisher, the rank and the p-value are listed.

**Supplementary Table 14. MAPR GSEA Annotations Specific to BTNR.**

Association tests with Gene Ontology terms were performed for the BTNR gene set using GSEA on the full gene ranking produced by MAPR for the gene set, "MAPR GSEA". The table showsthe Gene Ontology terms ("GO Term") that were in the top 50 enriched terms using the MAPR GSEA on BTNR "MAPR GSEA Rank", but the median rank of the term across the 9 related cancer gene sets ("Med Rank 9Sets") was worse than 100. Only GO terms of size ("SetSize") between 10 and 500 genes were considered. The last five columns show the percentage genes from the original BTRN gene set, the top 400 MAPR ranked genes, and the three protein families that are annotated for the given Gene Ontology term.

### Supplementary References

1. Warde-Farley, D., Donaldson, S.L., Comes, O., Zuberi, K., Badrawi, R., Chao, P., Franz, M., Grouios, C., Kazi, F., Lopes, C.T. *et al.* (2010) The GeneMANIA prediction server: biological network integration for gene prioritization and predicting gene function. *Nucleic Acids Res*, **38**, W214-220.
2. Liberzon, A., Subramanian, A., Pinchback, R., Thorvaldsdottir, H., Tamayo, P. and Mesirov, J.P. (2011) Molecular signatures database (MSigDB) 3.0. *Bioinformatics*, **27**, 1739-1740.
3. Goldman, M., Craft, B., Swatloski, T., Cline, M., Morozova, O., Diekhans, M., Haussler, D. and Zhu, J. (2015) The UCSC Cancer Genomics Browser: update 2015. *Nucleic Acids Res*, **43**, D812-817.
4. Kauppila, S., Stenback, F., Risteli, J., Jukkola, A. and Risteli, L. (1998) Aberrant type I and type III collagen gene expression in human breast cancer in vivo. *J Pathol*, **186**, 262-268.
5. Freire, J., Dominguez-Hormaetxe, S., Pereda, S., De Juan, A., Vega, A., Simon, L. and Gomez-Roman, J. (2014) Collagen, type XI, alpha 1: an accurate marker for differential diagnosis of breast carcinoma invasiveness in core needle biopsies. *Pathol Res Pract*, **210**, 879-884.
6. Raglow, Z. and Thomas, S.M. (2015) Tumor matrix protein collagen XIalpha1 in cancer. *Cancer Lett*, **357**, 448-453.
